# NDRG1 expression is an independent prognostic factor in inflammatory breast cancer

**DOI:** 10.1101/2020.09.25.313817

**Authors:** Emilly S Villodre, Yun Gong, Lei Huo, Esther C Yoon, Naoto T Ueno, Wendy A Woodward, Debu Tripathy, Juhee Song, Bisrat G Debeb

## Abstract

NDRG1 is widely described as a metastasis suppressor in breast cancer. However, we found that NDRG1 is critical in promoting tumorigenesis and brain metastasis in mouse models of inflammatory breast cancer (IBC), a rare but highly aggressive form of breast cancer. We hypothesized that NDRG1 is a prognostic marker associated with poor outcome in patients with IBC. Microarray gene expression data from the IBC Consortium dataset were analyzed to compare *NDRG1* expression between IBC and non-IBC tumors and among breast cancer subtypes. NDRG1 levels in tissue microarrays from 64 IBC patients were evaluated by immunohistochemical staining with anti-NDRG1 primary antibody (32 NDRG1-low [≤ median], 32 NDRG1-high [>median]). Overall and disease-free survival (OS and DSS) were analyzed with Kaplan–Meier curves and log-rank test. *NDRG1* mRNA expression was higher in IBC than in non-IBC tumors (*p*=0.007), and in more aggressive HER2+ and basal-like vs luminal IBC subtypes (*p*<0.0001). Univariate analysis showed NDRG1 expression, tumor grade, disease stage, estrogen receptor (ER) status, and receipt of adjuvant radiation to be associated with OS and DSS. NDRG1-high patients had poorer 10-year OS and DSS than NDRG1-low patients (OS, 19% vs 45%, *p*=0.0278; DSS, 22% vs 52%, *p*=0.0139). On multivariable analysis, NDRG1 independently predicted OS (hazard ratio [HR]=2.034, *p*=0.0274) and DSS (HR=2.287, *p*=0.0174). NDRG1-high ER-negative tumors had worse outcomes OS, *p*=0.0003; DSS, *p*=0.0003; and NDRG1-high tumors that received adjuvant radiation treatment had poor outcomes (OS, *p*=0.0088; DSS, *p*=0.0093). NDRG1 correlated positively with aggressive tumor characteristics in IBC and was a significant independent prognostic factor for DSS and OSS in IBC patients. Targeting NDRG1 may represent a novel strategy for improving clinical outcomes for patients with IBC.

---

Inflammatory breast cancer (IBC) is one of the most aggressive forms of breast cancer. Although rare, accounting for only 1%-4% of newly diagnosed breast cancer cases, it is responsible for a disproportionately high 10% of breast cancer-related deaths in the United States^1,2^. IBC has a unique biology characterized by rapid proliferation and metastasis; indeed, almost all patients have lymph node involvement and more than 33% of patients with IBC present with distant metastasis at the time of diagnosis ^3,4^. Even with multimodality treatment approaches that include systemic chemotherapy, surgery, and radiation therapy, prognosis for patients with IBC is worse than for non-IBC patients (overall survival [OS] rates 40% versus 63% at 5 years)^5-7^. This may be due in part to 70% of IBC patients presenting with aggressive subtypes of HER2+ or triple-negative breast cancer (TNBC), compared with 40% of non-IBC tumors^8^. Efforts have been undertaken to identify molecular markers and therapeutic targets distinct to IBC and have identified important targets and pathways including EGFR, E-cadherin, eIFG4I, RhoC, and TIG1/AXL^9-13^. However, no IBC-specific molecular signature or target has been identified thus far, and effective targeted therapies for this disease remain limited.

N-myc downstream regulated gene 1 (NDRG1) is a stress response protein involved in hypoxia, cell growth, lipid metabolism, and resistance to chemotherapy^14-19^. NDRG1 is widely known as a metastasis suppressor in breast cancer, acting mainly by suppressing migration and invasion of breast cancer cells^20-22^. However, we and others have shown NDRG1 to be a tumor promoter in aggressive breast cancer^23-25^. Nagai and colleagues also showed that high expression of NDRG1 was associated with aggressive breast cancer behaviors, including advanced stage at presentation and high-grade tumors and that NDRG1 was independently associated with poor survival outcome^26^. However, the expression of NDRG1 and its clinical importance in IBC remains unknown.

Herein, we examined the expression of NDRG1 by using immunohistochemical staining of a tissue microarray (TMA) composed of samples from IBC patients and evaluated the expression of NDRG1 and its correlation with survival outcomes. We also assessed the association between NDRG1 expression and outcome stratified by known prognostic factors. Our findings showed that high expression of NDRG1 in IBC tumors was an independent predictor of worse OS and disease-specific survival (DSS).

## RESULTS

The IBC Consortium dataset consists of microarray gene expression profiles of IBC and non-IBC tumors from three institutions^27^. With this dataset, we analyzed *NDRG1* expression in patients with IBC and non-IBC and observed that *NDRG1* was expressed at higher levels in patients with IBC relative to non-IBC (*p*=0.0287; Supplementary Figure 1a). Considering the breast cancer subtypes within IBC tumors, *NDRG1* expression was significantly increased in more aggressive basal-like and HER2+ subtypes than in luminal subtypes of IBC (*p*<0.0001; Supplementary Figure 1b). We further found that *NDRG1* expression was significantly higher in patients with TNBC versus non-TNBC tumors in both IBC and non-IBC patients (Supplementary Figure 1c), supporting that NDRG1 is positively associated with aggressive tumor characteristics. To determine whether NDRG1 protein expression is associated with outcome in IBC, immunohistochemical staining was performed on TMAs from 64 patients with primary IBC who were treated between 1991 and 2004 at The University of Texas MD Anderson Cancer Center. The average age of these patients was 50 years (range 23-75 years). Eighty three percent of patients were stage III, 80% high grade and 62% were ER-negative tumors, and 67% of these patients received adjuvant radiation. The median follow-up time for the patients studied was 11.7 years and the median OS time was 3.7 years. NDRG1 staining was predominantly cytoplasmic/membranous. Representative images of NDRG1-low and -high tumors are shown in Figure 1.

**Figure 1.**
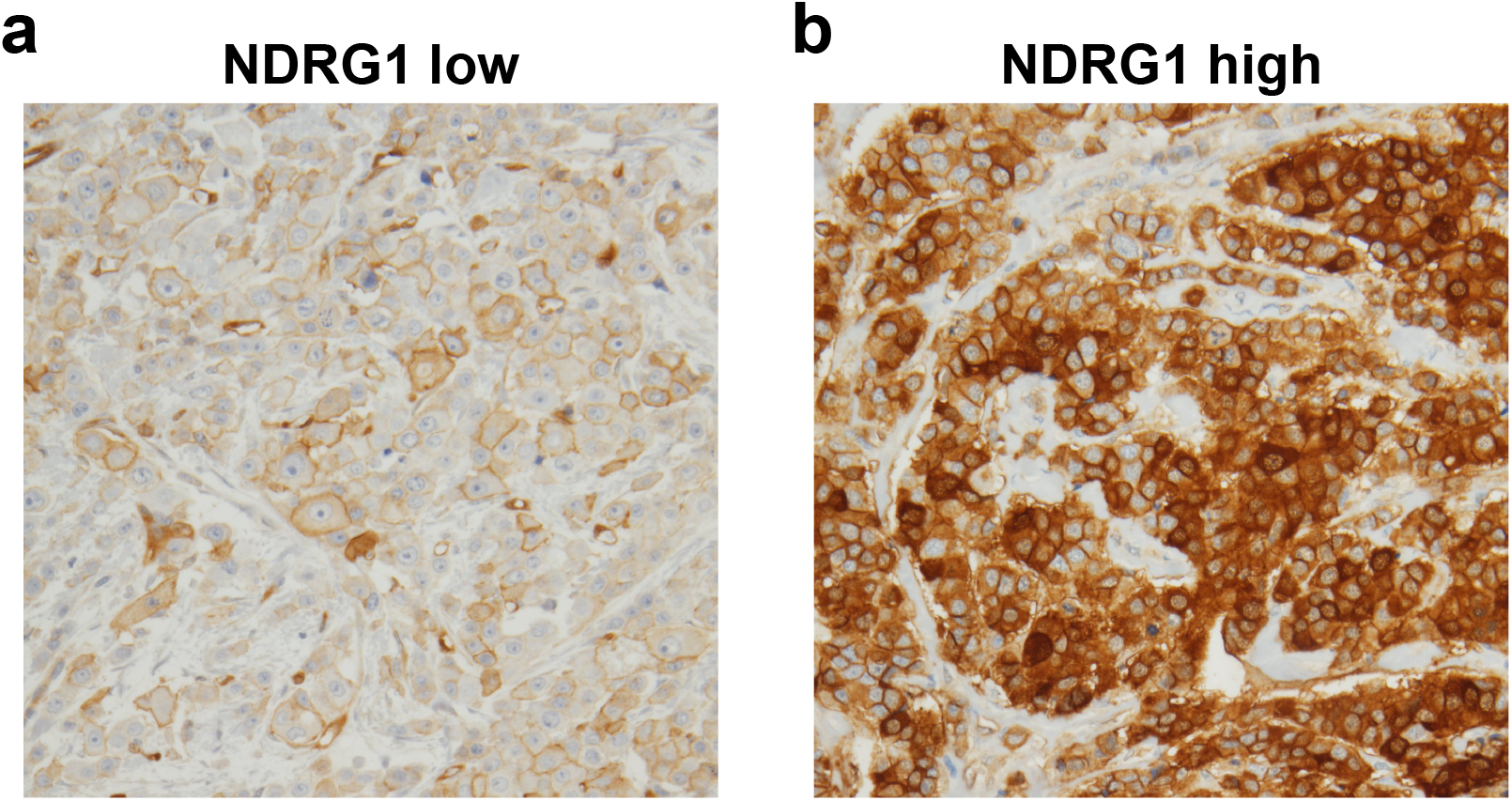
Immunohistochemical staining of NDRG1 in IBC tumors. Representative images of NDRG1 immunostaining of **(a)** an NDRG1-low IBC tumor and **(b)** an NDRG1-high IBC tumor.

Table 1 summarizes patient characteristics based on NDRG1 expression status, NDRG1 expression was associated with negative HER2 status (*p*=0.0077). Univariate analysis (Table 2) showed that NDRG1 expression (hazard ratio [HR]=2.1, *p*=0.0150), tumor grade (HR=2.4, *p*=0.0463), disease stage (HR=4.6, p=0.0011), hormone therapy (HR=0.3, *p*=0.0026), ER status (HR=0.4, *p*=0.0098), and adjuvant radiation therapy (HR=0.538, *p*=0.0434) were associated with OS. The same variables were also associated with DSS (Table 2).

**Table 1.** Clinical and pathologic characteristics of tumor samples from patients with IBC according to NDRG1 expression.

| **Covariate** | **Level** | **NDRG1-low (n=32)** | **NDRG1-high (n=32)** | **p-value** |
| --- | --- | --- | --- | --- |
| **Age** |  | 51.5 ± 12.1 | 48.6 ± 12 | 0.3861 |
| **Race** | Non-White | 8 (25%) | 5 (16%) | 0.5356 |
|  | White | 24 (75%) | 27 (84%) |  |
| **Histologic type** | Others | 2 (6%) | 4 (13%) | 0.6719 |
|  | Ductal | 30 (94%) | 28 (87%) |  |
| **Grade** | 1-2 | 6 (19%) | 7 (22%) | 0.7560 |
|  | 3 | 26 (81%) | 25 (78%) |  |
| **Lymphovascular invasion** | No | 4 (14%) | 3 (10%) | 0.7065 |
|  | Yes | 25 (86%) | 27 (90%) |  |
| **Stage** | III | 31 (97%) | 27 (84%) | 0.1961 |
|  | IV | 1 (3%) | 5 (16%) |  |
| **Estrogen receptor** | No | 21 (66%) | 19 (61%) | 0.7209 |
|  | Yes | 11 (34%) | 12 (39%) |  |
| **Progesterone receptor** | No | 24 (75%) | 19 (61%) | 0.2425 |
|  | Yes | 8 (25%) | 12 (39%) |  |
| **HER2** | No | 12 (37%) | 22 (71%) | **0.0077** |
|  | Yes | 20 (63%) | 9 (29%) |  |
| **Triple-negative breast cancer** | No | 27 (84%) | 20 (64%) | 0.0879 |
|  | Yes | 5 (16%) | 11 (36%) |  |
| **Adjuvant radiation** | No | 10 (31%) | 11 (34%) | 0.7901 |
|  | Yes | 22 (69%) | 21 (66%) |  |

**Table 2.** Univariate Cox regression analysis on overall survival and disease-specific survival among patients with IBC.

| Covariate | Level | Overall survival |  |  | Disease-specific survival |  |  |
| --- | --- | --- | --- | --- | --- | --- | --- |
|  |  | HR | 95% CI | p-value | HR | 95% CI | p-value |
| **Age** | 1 Unit Change | 1.007 | 0.980 - 1.035 | 0.6080 | 1.000 | 0.971 - 1.029 | 0.9904 |
| **NDRG1** | Low | 1.000 |  |  | 1.000 |  |  |
|  | High | 2.107 | 1.155 - 3.842 | **0.0150** | 2.354 | 1.235 - 4.485 | **0.0092** |
| **Race** | Non-white | 1.000 |  |  | 1.000 |  |  |
|  | White | 1.823 | 0.769 - 4.321 | 0.1724 | 1.940 | 0.757 - 4.973 | 0.1676 |
| **Histologic type** | Others | 1.000 |  |  | 1.000 |  |  |
|  | Ductal | 1.027 | 0.368 - 2.870 | 0.9591 | 1.153 | 0.410 - 3.243 | 0.7874 |
| **Grade** | 1-2 | 1.000 |  |  | 1.000 |  |  |
|  | 3 | 2.404 | 1.014 - 5.698 | **0.0463** | 2.612 | 1.020 - 6.688 | **0.0453** |
| **Lymphovascular invasion** | No | 1.000 |  |  | 1.000 |  |  |
|  | Yes | 1.684 | 0.601 - 4.717 | 0.3216 | 1.493 | 0.529 - 4.213 | 0.4486 |
| **Stage** | III | 1.000 |  |  | 1.000 |  |  |
|  | IV | 4.638 | 1.847 - 11.647 | **0.0011** | 5.485 | 2.138 - 14.069 | **0.0004** |
| **Estrogen receptor** | No | 1.000 |  |  | 1.000 |  |  |
|  | Yes | 0.414 | 0.212 - 0.808 | **0.0098** | 0.426 | 0.211 - 0.860 | **0.0173** |
| **Progesterone receptor** | No | 1.000 |  |  | 1.000 |  |  |
|  | Yes | 0.699 | 0.359 - 1.361 | 0.2919 | 0.816 | 0.412 - 1.616 | 0.5598 |
| **HER2** | No | 1.000 |  |  | 1.000 |  |  |
|  | Yes | 0.748 | 0.409 - 1.366 | 0.3445 | 0.602 | 0.313 - 1.161 | 0.1300 |
| **Triple-negative breast cancer** | No | 1.000 |  |  | 1.000 |  |  |
|  | Yes | 1.557 | 0.810 - 2.992 | 0.1844 | 1.692 | 0.852 - 3.358 | 0.1328 |
| **Adjuvant radiation** | No | 1.000 |  |  | 1.000 |  |  |
|  | Yes | 0.538 | 0.295 - 0.982 | **0.0434** | 0.486 | 0.259 - 0.914 | **0.0252** |

The Kaplan-Meier method was used to evaluate the association of NDRG1 expression and survival over time. Patients with NDRG1-low tumors experienced better actuarial 10-year OS (*p*=0.0129, Figure 2a) and DSS (*p*=0.0134, Figure 2b). Patients with NDRG1-high tumors showed significantly lower 10-year OS and DSS rates than patients with NDRG1-low (OS, 19% vs. 45%, *p*=0.0278; DSS, 22% vs 52%, *p*=0.0139). The median OS and DSS times were shorter for NDRG1-high patients (OS 2.5 years; DSS, 3.1 years) than for NDRG1-low patients (OS, 5.9 years; DSS, 10.7 years). Multivariable model predictors of OS and DSS included NDRG1 expression, ER status, disease stage, and receipt of adjuvant radiation (Table 3). Tumor grade was identified as being associated with OS and DSS at the univariate level but not at the multivariable level. NDRG1-high expression was a strong independent predictor of OS (HR=2.034, 95% confidence interval [CI]=1.081–3.822, *p*=0.0274) and DSS (HR=2.287, 95% CI=1.157–4.522, *p*=0.0174).

**Table 3.** Multivariate Cox regression analysis on overall survival and disease-specific survival among patients with IBC.

| Covariate | Level | Overall survival |  |  | Disease-specific survival |  |  |
| --- | --- | --- | --- | --- | --- | --- | --- |
|  |  | HR | 95% CI | p-value | HR | 95% CI | p-value |
| **NDRG1** | Low | 1.000 |  |  | 1.000 |  |  |
|  | High | 2.034 | 1.081 - 3.822 | **0.0274** | 2.287 | 1.157 - 4.522 | **0.0174** |
| **Estrogen receptor** | No | 1.000 |  |  | 1.000 |  |  |
|  | Yes | 0.313 | 0.156 - 0.628 | **0.0011** | 0.307 | 0.147 - 0.642 | **0.0017** |
| **Stage** | III | 1.000 |  |  | 1.000 |  |  |
|  | IV | 3.212 | 1.171 - 8.814 | **0.0234** | 3.844 | 1.362 - 10.854 | **0.0110** |
| **Adjuvant radiation** | No | 1.000 |  |  | 1.000 |  |  |
|  | Yes | 0.562 | 0.298 - 1.058 | 0.0741 | 0.510 | 0.261 - 0.994 | **0.0481** |

**Figure 2.**
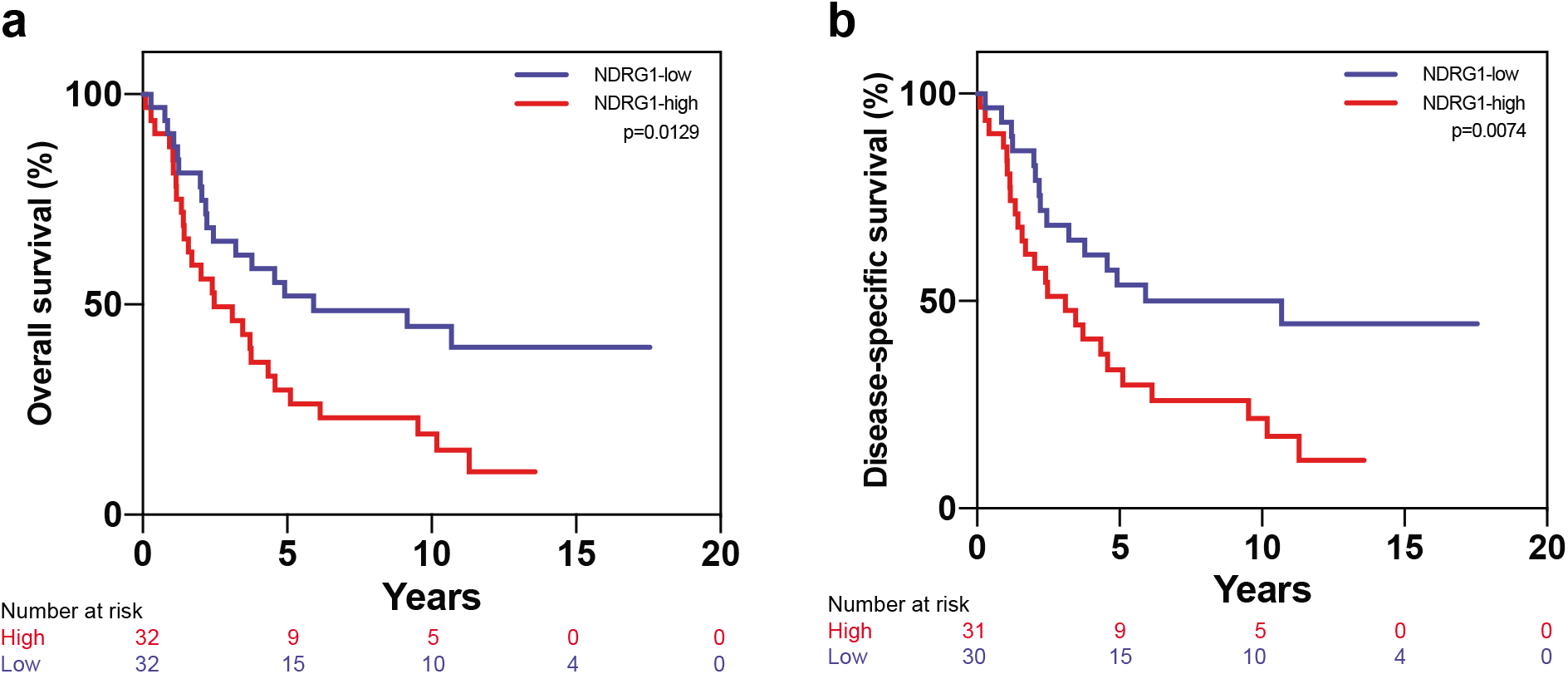
NDRG1 is a predictor of poor outcome in patients with IBC. Kaplan-Meier analysis showed that patients whose tumors had NDRG1-high expression had **(a)** worse overall survival and **(b)** worse disease-specific survival than did patients whose tumors had NDRG1-low expression.

ER status was also an important prognostic factor for OS and DSS for patients with IBC; patients with ER-negative tumors had worse OS (*p*=0.0077) and DSS (*p*=0.01) relative to patients with ER-positive IBC tumors (Figure 3a-b). Multivariable analysis showed ER status to be an independent factor associated with OS (HR=0.307, 95% CI=0.145–0.650, *p*=0.002) and DSS (HR=0.307, 95% CI=0.147–0.642, *p*=0.0017) (Table 3). Interestingly, NDRG1-high and ER-negative tumors were associated with the worst clinical outcomes for patients with IBC (OS, *p*=0.0003; DSS, *p*=0.0003; Figure 3c-d). Survival outcomes of ER-positive patients were not affected by NDRG1 expression (Figure 3c-d). Analysis of median OS and DSS times highlights the importance of stratifying patients for both variables: patients with ER-negative tumors had a median 2.2 years for both OS and DSS, whereas those with ER-negative / NDRG1-high tumors had a median 1.6 years, and ER-negative / NDRG1-low tumors had medians of 3.2 years OS and 4.6 years DSS (Figure 3e-f).

**Figure 3.**
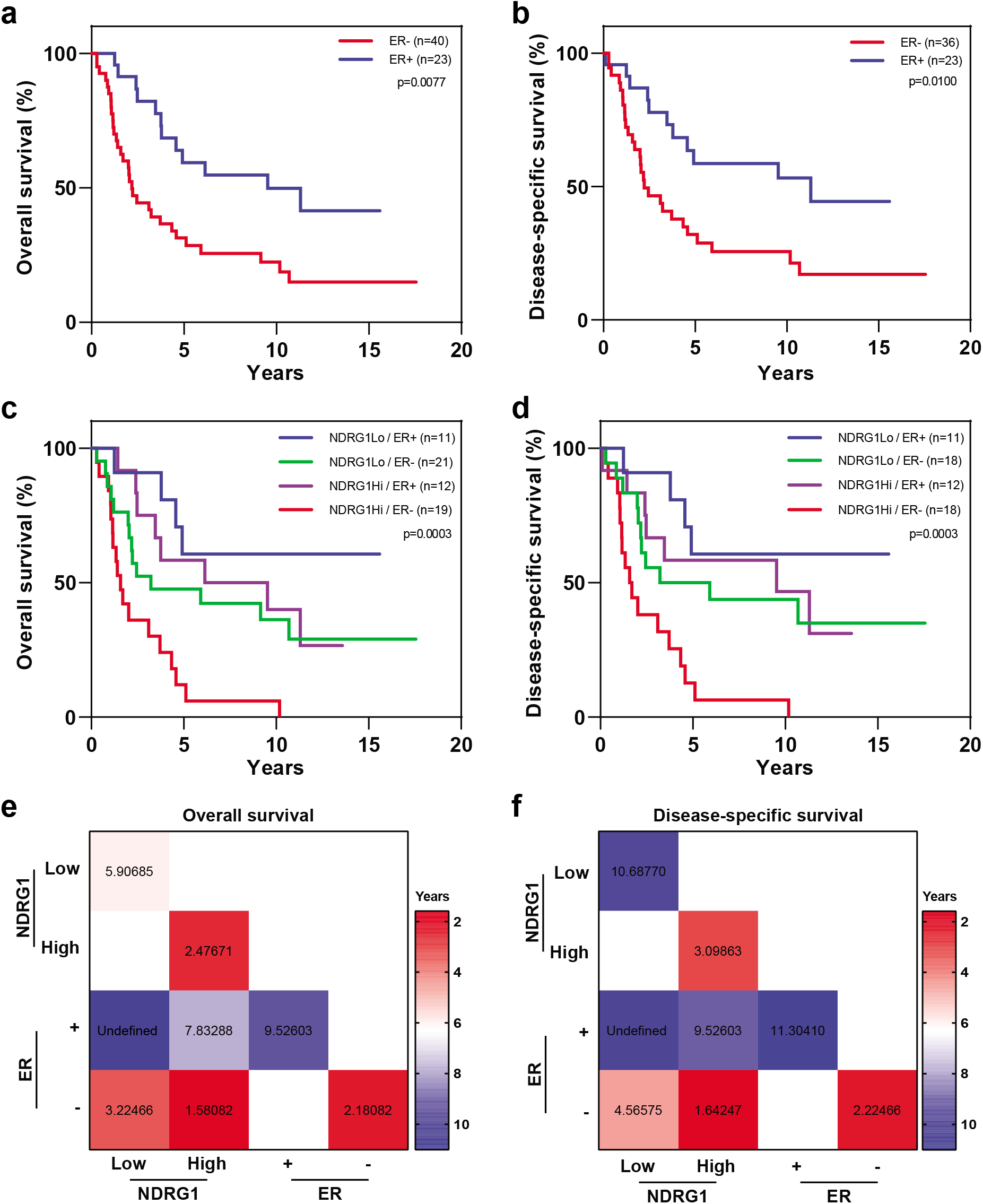
Overall survival and disease-specific survival in patients with IBC stratified by ER status and NDRG1 expression. Patients with estrogen receptor (ER)-negative tumors had **(a)** worse overall survival and **(b)** worse disease-specific survival versus patients with ER-positive tumors. **(c, d)** Stratification of patients by ER and NDRG1 expression status in terms of overall survival and diseasespecific survival. Log-rank tests were used to obtain *p* values. **(e, f)** Median overall survival and disease-specific survival times, in years, for patients stratified by ER status and NDRG1 expression.

Disease stage was another prognostic variable for OS (HR=3.212, 95% CI=1.171–8.814, *p*=0.0234) and DSS (HR=3.844, 95% CI=1.362–10.854, *p*=0.011). Kaplan-Meyer analysis showed that patients with stage III IBC had better outcomes than did patients with stage IV tumors (OS, *p*=0.0003; DSS, *p*<0.0001) (Figure 4a-b). Further stratification of patients with stage III disease according to NDRG1 expression status showed a significant difference in outcomes, wherein patients with stage III NDRG1-high tumors had worse OS (*p*=0.045) and DSS (*p*=0.0239) than did patients with stage III NDRG1-low tumors (Figure 4c-d). We could not perform similar analyses of stage IV tumors owing to small patient numbers. Interestingly, the median OS times for patients with stage III tumors differed considerably by NDRG1 expression level, being 9.1 years in NDRG1-low tumors to 4.6 years for NDRG1-high stage III tumors (Figure 4e). Similarly, the median DSS times were 4.9 years for stage III NDRG1-low tumors and 10.7 years for stage III NDRG1-high tumors (Figure 4f).

**Figure 4.**
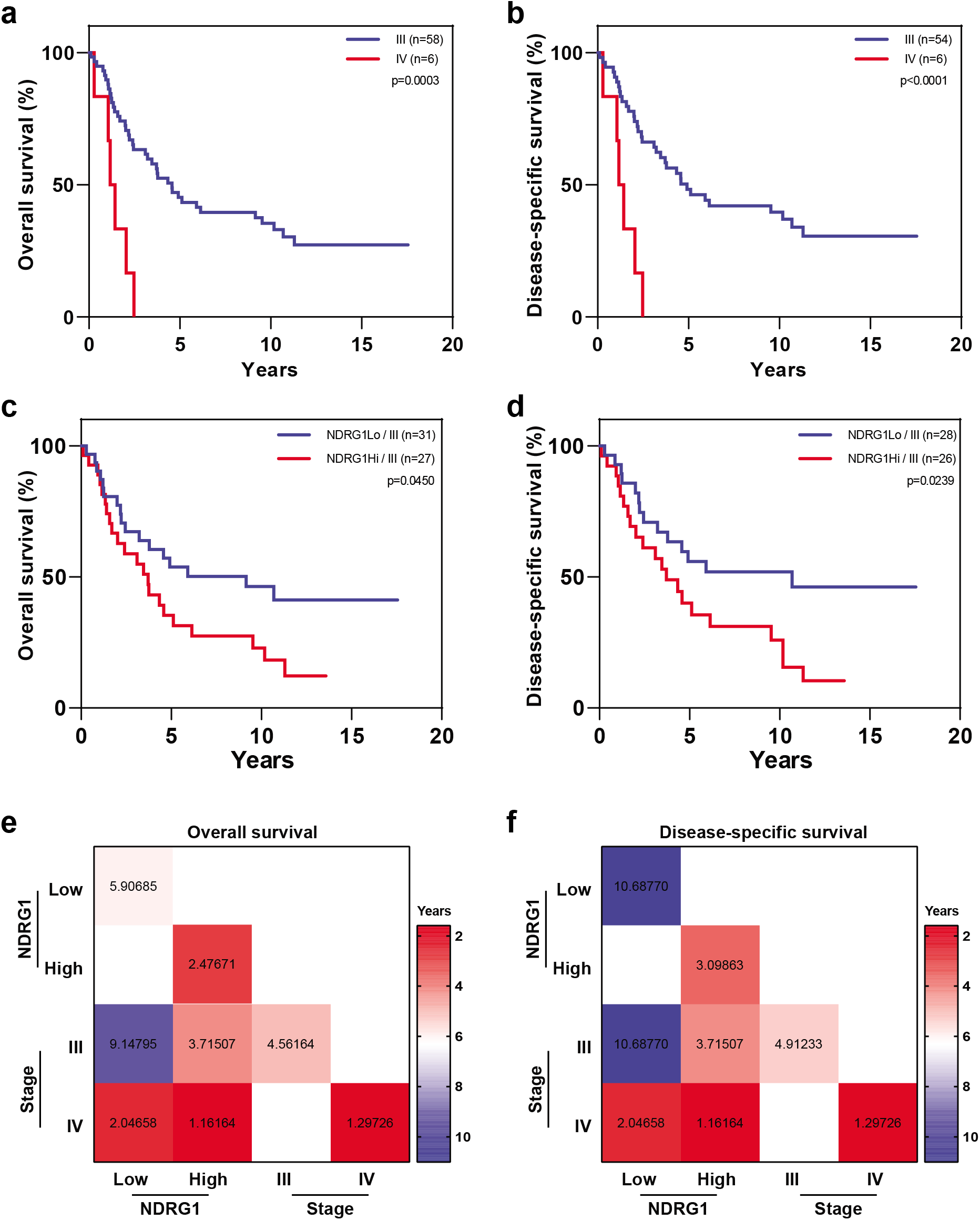
Overall survival and disease-specific survival in patients with IBC stratified by disease stage and NDRG1 expression. Patients with stage III IBC had better **(a)** overall survival and **(b)** disease-specific survival than did patients with stage IV IBC. **(c, d)** Patients with stage III IBC stratified by NDRG1 expression in terms of overall survival and disease-specific survival. Log-rank tests were used to obtain *p* values. **(e, f)** Median overall survival and disease-specific survival times, in years, for patients stratified by disease stage and NDRG1 expression.

Receipt of adjuvant radiation was also an independent variable related to DSS (HR=0.510, 95% CI=0.261–0.994, *p*=0.0481) (Table 3). Patients who received adjuvant radiation had better survival outcomes than those who did not (OS, *p*=0.0403; DSS, *p*=0.0137) (Figure 5a-b). Among patients who received adjuvant radiation, those with NDRG1-high tumors showed poorer outcomes than those with NDRG1-low tumors (OS, *p*=0.0088; DSS, *p*=0.0128, Figure 5c-d). Among patients who did not receive adjuvant radiation therapy, NDRG1 expression did not correlate with survival outcomes (Figure 5e-f). The median survival times for all patients who received adjuvant radiation was 3.7 years for OS and 4.6 years for DSS. Stratification of radiation-treated patients by NDRG1 again showed distinct differences in survival time, with medians of 3.1 years for both OS and DSS for NDRG1-high tumors versus not achieved for NDRG1-low tumors (Figure 5g-h).

**Figure 5.**
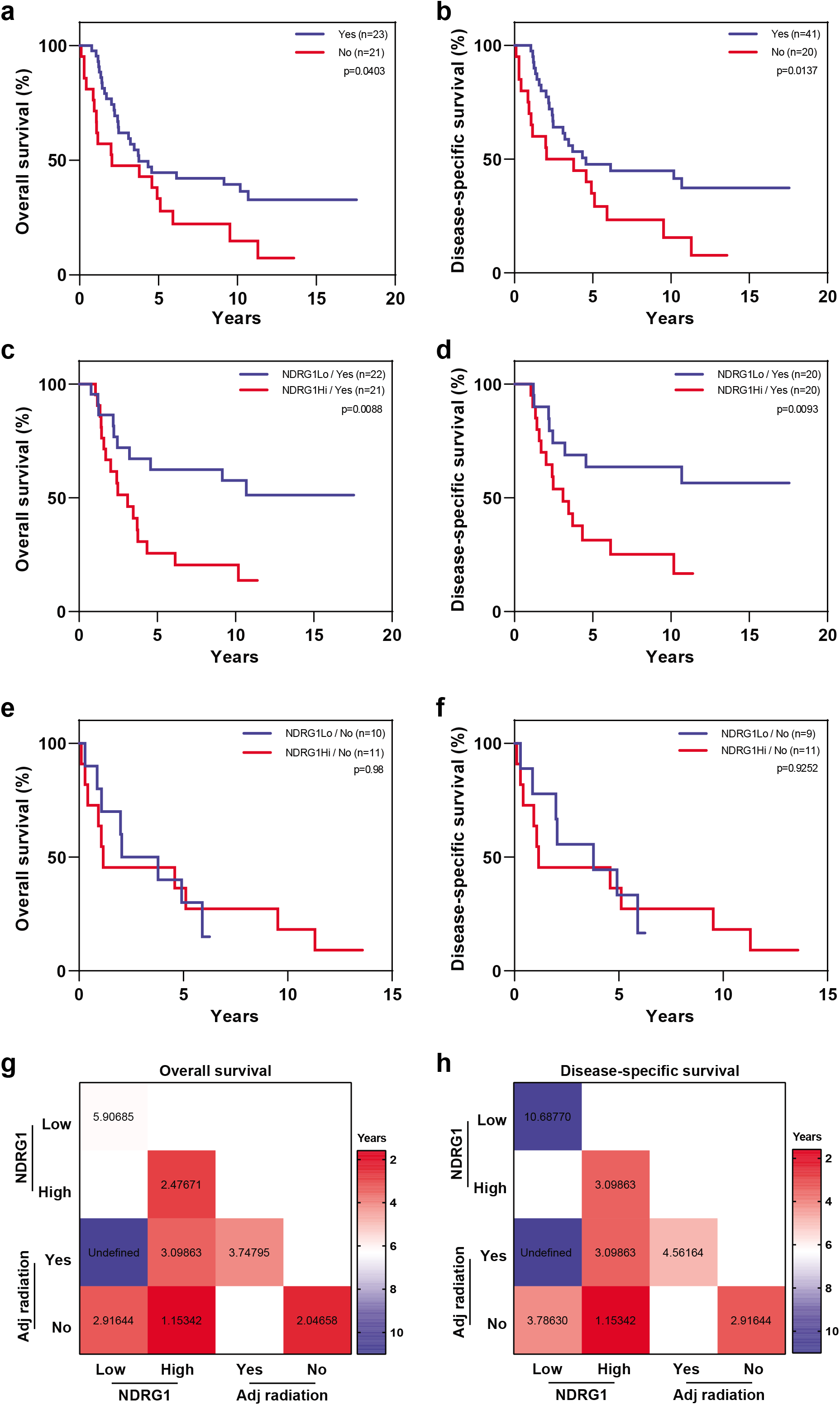
High NDRG1 expression correlated with worse outcomes among patients who received adjuvant radiation therapy. IBC patients who received adjuvant radiation had better **(a)** overall survival **(b)** and disease-specific survival than did patients who did not receive radiation. **(c, d)** Patients who received adjuvant radiation treatment stratified by NDRG1 expression in terms of overall survival and disease-specific survival. **(e, f)** Patients who did not receive adjuvant radiation stratified by NDRG1 expression in terms of overall survival and disease-specific survival. Log-rank tests were used to obtain *p* values. **(g, h)** Median overall survival and disease-specific survival times, in years, for patients stratified by NDRG1 expression and adjuvant radiation treatment status.

As expected, patients with lower tumor grades (I-II) had better outcomes than those with high-grade tumors (OS, *p*=0.0399; DSS, *p*=0.0386; Figure 6a-b). Despite the small number of low-grade tumors, we observed a significant difference in OS (*p*=0.0363) and DSS (*p*=0.0210) after stratifying for NDRG1-high versus NDRG1-low expression; patients with low-grade tumors and NDRG1-low expression had better outcomes than patients with NDRG1-high expression (Figure 6c-d). Outcomes may have been worse for patients with high-grade tumors and NDRG1-high expression relative to those with NDRG1-low expression, but those apparent differences were not statistically significant (OS, *p*=0.0765; DSS, *p*=0.0699) (Figure 6d-e). Low-grade, NDRG1-high tumors were associated with shorter survival, with median survival times of 4.3 years versus not achieved for low-grade, NDRG1-low tumors (Figure 6g-h).

**Figure 6.**
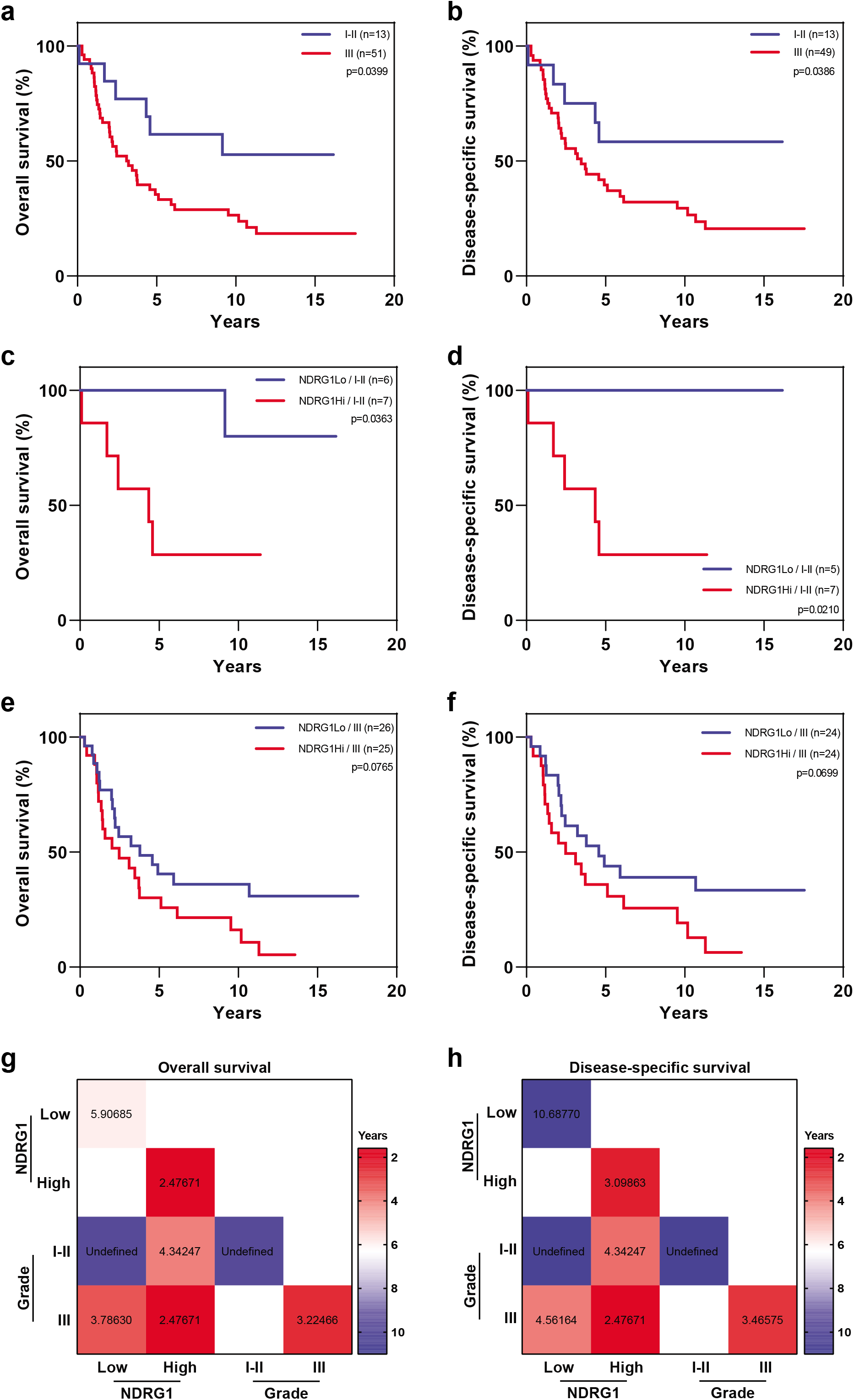
Overall survival and disease-specific survival in patients with IBC stratified by tumor grade and NDRG1 expression. Patients with IBC and low-grade cancer had better **(a)** overall survival and **(b)** disease-specific survival than did patients with grade III disease. **(c, d)** Patients with low-grade IBC stratified by NDRG1 expression in terms of overall survival and disease-specific survival. **(e, f)** Patients with high-grade IBC stratified by NDRG1 expression in terms of overall survival and disease-specific survival. Log-rank tests were used to obtain *p* values. **(g, h)** Median overall survival and disease-specific survival times, in years, for patients stratified by NDRG1 expression and tumor grade.

## DISCUSSION

IBC remains a relatively poorly defined disease that lacks specific therapeutic targets and prognostic biomarkers; the molecular characterization of IBC could advance our understanding of its unique biology and provide opportunities that could be translated into novel therapeutic strategies to improve clinical outcomes. Herein, we report that NDRG1 protein expression was an independent predictor of poor survival outcomes for patients with IBC. In subset analyses, we report that NDRG1-high expression in patients with ER-negative, stage III tumors, and patients who received adjuvant radiation had worse outcomes than did patients with NDRG1-low tumors. Our results suggest that IBC patients could be stratified not only by known prognostic markers but also by biological determinants such as NDRG1 expression status. Further, we found that *NDRG1* mRNA was overexpressed significantly in IBC relative to non-IBC, and in HER2+, basal-like, and TNBC aggressive breast cancer subtypes compared with luminal-like IBC tumors. These findings support our contention that NDRG1 is associated with aggressive tumor features in IBC.

*NDRG1* is a stress response gene that is highly activated and expressed in hypoxia and resistance to chemotherapy. Its function in breast cancer is widely described as a tumor and metastasis suppressor, acting mainly through inhibition of migration and invasion of cancer cells^20-22,28^. The induction of *NDRG1* was shown in a mouse mammary tumor model to suppress metastasis by modulating WNT pathway signaling^21^. Chiang *et al.* also described how silencing *NDRG1* expression in MCF-7 cells led to increased proliferation and invasiveness of those breast cancer cells^28^. Paradoxically, other studies showed that *NDRG1* may function as an oncogene or a prognostic biomarker in aggressive forms of breast cancer^19,23,24,26^. Mao *et al.* found that NDRG1 could be used as a marker for invasive breast cancer, observing that NDRG1 expression was significantly higher in invasive breast cancer versus matched non-tumor tissues, and its levels were associated with progression from breast atypia to carcinoma. They also observed a correlation between advanced tumor stage and high NDRG1 expression^23^. Nagai *et al.* found associations between high NDRG1 expression and worse DSS and OS in a cohort of 600 patients: the 10-year OS rate was 35% for NDRG1-high versus 67% for NDRG1-low, and NDRG1 was an independent prognostic factor for both OS and DSS. Moreover, NDRG1 was expressed at higher levels in stage III and IV breast cancer and in grade 3 tumors^26^. More recently, a study by Sevinsky and colleagues demonstrated that NDRG1 promotes breast cancer aggressiveness by altering lipid metabolism^19^.These observations are supported by our previous work showing that NDRG1 promotes tumorigenesis and brain metastasis in mouse models of aggressive breast cancer^25^.

Expression of the ER is a well-known prognostic and predictive factor, and ER status is essential in the choice of treatment strategy. Patients with ER-positive breast cancer benefit from the use of hormonal therapy and have better OS than do patients with ER-negative disease, and this improvement is independent of disease stage and tumor grade^29-31^. Our results in the present study confirmed that ER status was, indeed, an independent factor related to both OS and DSS, with ER-negative IBC patients exhibiting worse clinical outcomes. ER-negative status is associated with aggressive growth and shorter survival. Interestingly, in our study stratification of ER-negative patients by NDRG1 expression level showed significant differences in survival outcomes: ER-negative, NDRG1-high tumors were associated with worse outcomes than ER-negative, NDRG1-low tumors. However, no such difference was observed in ER-positive tumors stratified by NDRG1 expression. Our findings indicate that the clinical outcome of patients with ER-negative IBC can be stratified further based on NDRG1 expression status.

Adjuvant radiation therapy is an important part of breast cancer treatment and is known to improve breast cancer-specific survival and reduce tumor recurrence^32-34^. In the current study, we found that receipt of adjuvant radiation for IBC tumors independently correlated with improved breast cancer-specific survival. We also showed that patients who received adjuvant radiation and had NDRG1 low-expressing tumors had better clinical outcomes than did those with NDRG1-high-expressing radiation-treated tumors. These hypothesis-generating findings suggest that the role of NDRG1 in local failure in breast cancer patients and radiation resistance warrants further investigation. Previous studies of rectal cancer cells have shown that NDRG1 is one of the top highly upregulated genes in response to ionizing radiation, and that depleting it could be a promising strategy to sensitize cells to radiotherapy^35^. Many studies have been conducted to develop a “radiosensitivity signature” to stratify patients according to benefit from adjuvant radiation treatment^36-38^. However, no such molecular signature for radiation response has yet been identified.

In conclusion, NDRG1 expression was an independent prognostic factor for worse survival outcomes in patients with IBC, and together with other important prognostic factors, such as ER status and disease stage, can be used to further stratify prognostic outcome or treatment response. Because NDRG1 expression levels are generally high in IBC, targeting NDRG1 may provide a novel therapeutic strategy to improve outcome for patients with IBC.

## MATERIALS AND METHODS

### IBC tumor microarrays and immunohistochemical staining

This study was approved by institutional review board of The University of Texas MD Anderson Cancer Center. Details of disease diagnosis, preoperative and postoperative treatments, biomarker studies (including ER, PR, and HER2 status), and TMA construction with post-neoadjuvant residual tumors are reported elsewhere^39^. Immunohistochemical staining of TMAs was done with a monoclonal antibody against NDRG1 (1:5000, #9485, Cell Signal) that was previously validated^40^. NDRG1 staining was evaluated by percentage (0%–100%) and intensity (weak, moderate and strong) of invasive tumor cells showing cytoplasmic and/or membranous staining. NDRG1 H-score was calculated by multiplying the percentage with intensity, and the NDRG1 H-score median was used as cutoff, wherein 32 patients were grouped as NDRG1-low (≤ median), and 32 as NDRG1-high (>median). Representative images of NDRG1-low and NDRG1-high tumors are shown in Figure 1.

### Analysis of IBC Consortium dataset

We analyzed microarray gene expression data from the multi-institutional IBC Consortium dataset, which contains 137 IBC and 252 non-IBC tumors, to compare *NDRG1* mRNA expression between IBC and non-IBC tumors and among the molecular subtypes of breast cancer^27^.

### Statistical Analysis

Patient characteristics were summarized by NDRG1 value (Low [≤median] vs. High [>median]) and compared between patients with NDRG1-low and patients with NDRG1-high tumors. Two-sample *t* tests or Wilcoxon rank-sum tests were used for the comparison of continuous variables. Chi-square tests or Fisher’s exact tests were used for the comparison of categorical variables. OS was defined as the interval from diagnosis to death, and DSS as the interval from diagnosis to death from breast cancer. Those patients without an event (death or breast cancer death) were censored at last follow-up. Kaplan-Meier curves and log-rank tests were used to compare survival distributions. Univariate and multivariate Cox proportional hazards regression models were used to compare OS (and DSS) between NDRG1-low and -high groups, adjusting for other covariates. The proportional hazards assumption was checked by scaled Schoenfeld residual plots and correlation between the scaled Schoenfeld residuals and survival time. *P* values of <0.05 indicated a statistically significant difference. SAS 9.4 (SAS institute INC, Cary, NC) was used for data analysis.

For analyses of the IBC Consortium dataset, patients were stratified as having NDRG1-high-expressing or NDRG1-low-expressing tumors according to the median *NDRG1* mRNA expression levels in the tumor samples. Mann-Whitney tests were used to compare the two categories. Black lines in each group indicate median ± SD. GraphPad software (GraphPad Prism 8, La Jolla, CA) was used. *P* values of <0.05 were considered to indicate significant differences.

## ACKNOWLEDGEMENTS

We thank Christine F. Wogan for scientific editing of the manuscript, and Carol M. Johnston from the Division of Surgery Histology Core at UT MD Anderson for her help with immunohistochemical staining.

## Funding

This study was supported in part by the following grants: Susan G. Komen Career Catalyst Research grant (CCR16377813 to BGD), American Cancer Society Research Scholar grant (RSG-19-126-01 to BGD), and the State of Texas Rare and Aggressive Breast Cancer Research Program.

## AUTHOR CONTRIBUTIONS

E.S.V. and B.G.D. conceived and designed the project. E.S.V. performed experiments, analyzed data and interpreted the results. Y.G. provided TMA of IBC patient samples. Y.G. L.H and E.C.Y. analyzed the TMA and interpreted results. J.S. provided statistical analysis support. N.T.U, W.A.W, and D.T. provided resources and contributed to revision of the manuscript. E.S.V. and B.G.D. wrote and edited the manuscript with input from all other authors.

## COMPETING INTERESTS STATEMENT

The authors declare no conflict of interest.

## Notes

### Competing Interest Statement

The authors have declared no competing interest.

