## Supplementary Figure 1 for "NDRG1 expression is an independent prognostic factor in inflammatory breast cancer"

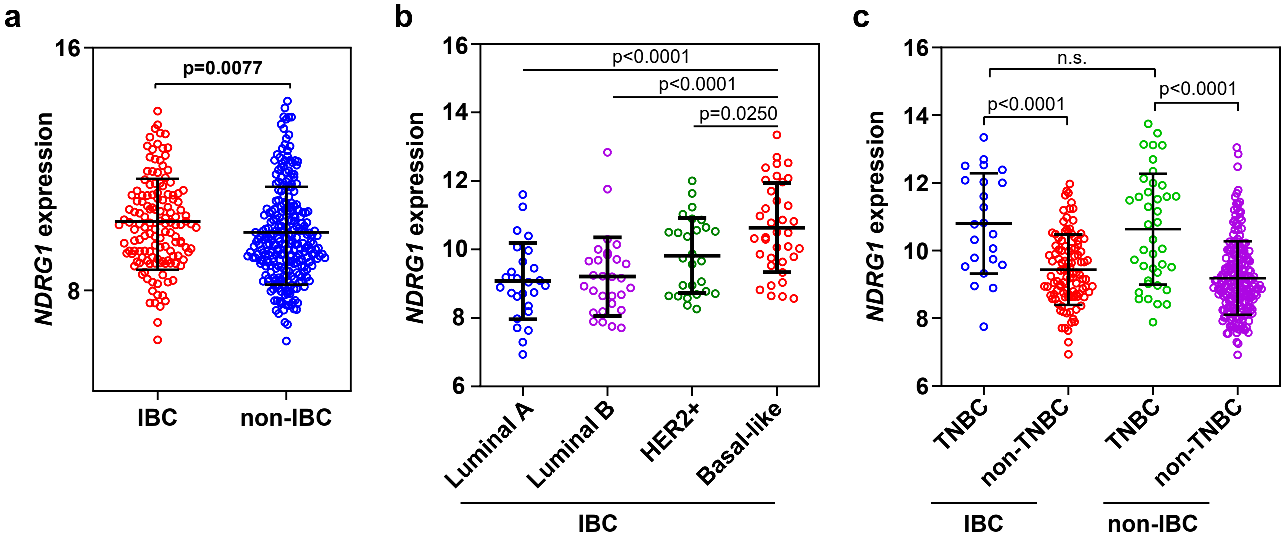


**Supplementary Figure 1. *NDRG1* is highly expressed in more aggressive molecular subtypes of IBC. (a)** IBC patients express higher *NDRG1* levels compared to non-IBC patients. **(b)** IBC patients with more aggressive molecular subtype (basal-like and HER2+) showed higher levels of *NDRG1* versus less aggressive, luminal molecular subtype. **(c)** *NDRG1* expression is higher in triple-negative (TNBC) compared to non-TNBC in both IBC and non-IBC patients. Black lines in each group indicate median ± SD. p values were calculated using Mann-Whitney test.
